# An inhibitory circuit motif governs oscillation-dependent coupling between aperiodic activity and neural spiking

**DOI:** 10.64898/2026.04.16.719083

**Authors:** Janna D. Helfrich, Julia Veit, Randolph F. Helfrich

## Abstract

Behavior arises from coordinated neural population activity, yet how spiking and synaptic interactions give rise to extracellular population signals, including local field potentials (LFPs), remains unresolved. Beyond oscillations, aperiodic LFP activity has emerged as a putative marker of neural excitability, but its neurophysiological basis and relationship to neural spiking are unclear. Here we employed optogenetics and simultaneous single-unit and LFP recordings in mouse visual cortex to quantify how spiking relates to oscillatory and aperiodic dynamics across cortical states and behavioral contexts. To test circuit mechanisms, we optogenetically suppressed the activity of somatostatin (SST), vasoactive intestinal peptide (VIP), or parvalbumin (PV) interneurons to causally manipulate inhibitory drive, thereby dissociating spiking, aperiodic activity, and gamma oscillations. Across conditions, aperiodic activity tracked neural spiking in a manner consistent with computational predictions, but this coupling was strongly context- and state-dependent: high oscillatory synchrony attenuated the relationship between aperiodic activity and spiking. These results demonstrate that state-dependent changes in neural excitability shift circuit activity between oscillation- and aperiodic-dominated regimes, placing principled limits on interpreting neural mass signals at synaptic or cellular scales.

## Introduction

Recordings of extracellular electric fields, such as local field potentials (LFPs), are widely used to study coordinated population activity that supports perception, action, and cognition^1–6^. Yet despite their widespread use, the precise mechanistic relationship between single neuron spiking and LFPs remains poorly understood^1,2,7,8^. Despite this gap, LFP activity is often interpreted as a window into coordinated population dynamics in different experimental scenarios, behavioral contexts, or pathological conditions^3–6^. Among the LFP features, gamma-band oscillations (∼30-80 Hz) have received particular attention and are thought to support information acquisition, integration, and inter-areal communication in large-scale cortical networks^3,6,9,10^. Substantial efforts were therefore devoted to characterizing the circuit dynamics that give rise to gamma oscillations, motivated by the idea that different oscillatory patterns reflect distinct cellular substrates and computational operations^9,11–13^. In canonical accounts, gamma emerges from the recurrent interactions between excitatory pyramidal cells and inhibitory interneurons, forming the ‘gamma cycle’ that provides a temporal reference frame for computation through coordinated interplay between excitation and inhibition^9,13,14^. Consistent with this framework, interneuron-mediated inhibition through parvalbumin- (PV), somatostatin-(SST) and vasoactive intestinal peptide-positive (VIP) interneurons have been identified as key organizers that connect cortical assemblies into functional units^15–24^.

Despite this broad interest, recent discoveries indicate that gamma oscillations are strongly stimulus- and context-dependent; limiting their presumed role for domain-general cognition^25–29^. In addition, narrow-band gamma oscillations are frequently conflated with broadband power increases that reflect the 1/f decay function of electrophysiological power spectra^25–27,30^. Accordingly, broadband high-gamma (∼80-150 Hz) can dissociate from narrow-band oscillatory gamma^25,27,31^ and neither component maps directly onto spiking activity^8,32–34^. Hence, the neurophysiological basis of broadband activity (sometimes also referred to as aperiodic or scale-free activity) and its relationship to spiking activity remain unclear. Computational modeling has suggested that the aperiodic 1/f slope of the power spectrum may reflect the balance of excitation and inhibition^35,36^, but this interpretation is debated on theoretical and empirical grounds^37–39^, including pharmacological^40–42^, electrophysiological and in-vivo imaging studies^43^. However, it remains a popular interpretation as broadband changes are ubiquitous in electric field recordings. Much of the existing evidence comes from modeling or human recordings, leaving open how inhibitory circuit mechanisms causally shape aperiodic activity in vivo. Here, we test how inhibitory drive controls aperiodic and oscillatory activity and their coupling to spiking. We performed simultaneous single-unit and LFP recordings in mouse V1^23,24^ during variations in cortical state (locomotion vs. quiescence) and visual context (different gratings vs. blank screen). To causally manipulate inhibition within a canonical circuit motif^44,45^, we optogenetically suppressed the activity of the three cardinal inhibitory interneuron-types (PV+, SST+, VIP+) in separate experiments (**Fig. 1a**), enabling joint analysis of neural spiking, synchrony, and population-level aperiodic activity across cortical states (**Fig. 1b/c**). Using spectral parametrization^46^ to dissociate oscillatory peaks from aperiodic background (**Fig. 1d-f**), we quantify how inhibitory microcircuit control governs neural excitability and shapes the LFP signatures that are commonly used as proxies for circuit computation.

**Figure 1.**
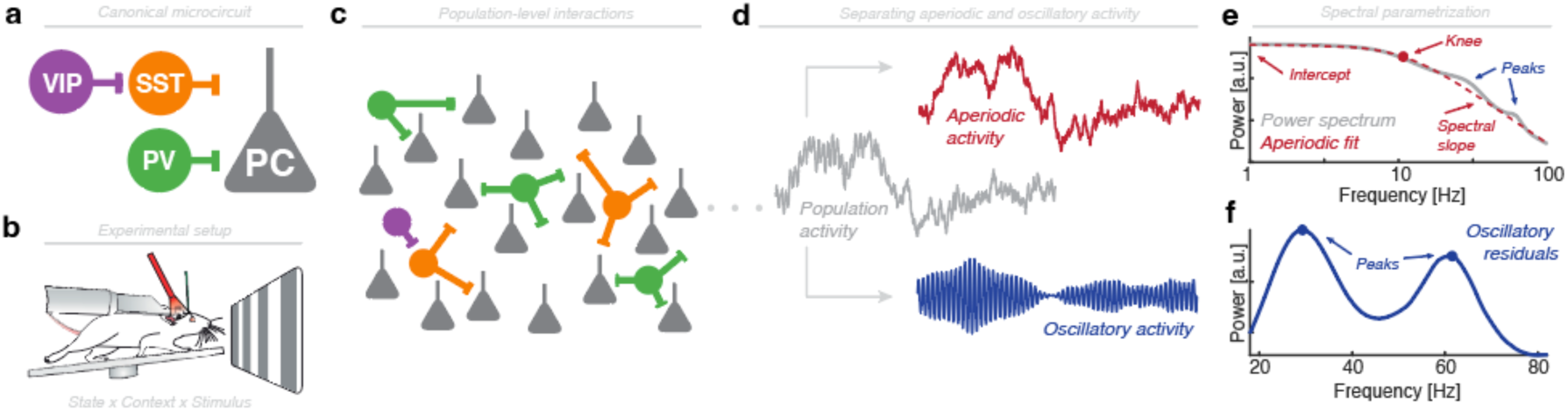
Circuit model, experimental and analytical approach. (**a**) Simplified microcircuit model describing the interactions between pyramidal cells (PC) and interneuron activity PV+ (green), SST+ (orange) and VIP+ (purple)-positive interneurons. Note that the interneuron subtypes have different suspected PC targets (direct/indirect; soma/dendrite). (**b**) Experimental setup: Mice were running or resting on a treadmill, while facing a screen that displayed gratings of varying sizes or remained blank, while different interneurons were optogenetically inactivated in different mice. (**c**) The neuronal population is composed of ∼80% PCs and ∼20% interneurons, where PV-positive interneurons reflect the largest subgroup. (**d**) Coordinated population spiking gives rise to the local field potential (in grey), which consists of aperiodic activity that follows a 1/f scale-free dynamics (red) and oscillatory activity (blue). (**e**) In the frequency domain, aperiodic activity can be estimated from the power spectrum (grey) and can be described by few parameters (offset, spectral knee and its exponent (the absolute value of the negative spectral slope); in red). Oscillations are evident as discrete peaks above the aperiodic fit (blue). (**f**) Oscillatory residuals (blue) can be obtained by subtracting the aperiodic fit from the power spectrum (cf. panel **e**) and can be described by their peak frequency and amplitude. Panel **b** is reproduced from23 with permission from the authors.

## Results

To determine how neural spiking relates to aperiodic and oscillatory activity, we analyzed neural activity across cortical states, visual stimulation contexts, and optogenetic suppression of interneuron subtypes (sub-selections based on availability; *Methods*) from 60 mice (SST-Cre, VIP-Cre and PV-Cre). LFP power spectra were parametrized to estimate the aperiodic component (*Methods*) and defined the oscillatory residuals as the difference between the original power spectrum and the aperiodic fit. Model goodness-of-fits were consistently high (99.2 ± 2.1%, median ± SD; **Supplementary Text** and **Fig. S1**). Subsequent analyses of neural oscillations focused on size-dependent visual gamma activity (∼25-40 Hz in mice) and did not target other oscillatory phenomena, such as locomotion-related theta or fast gamma activity at ∼60 Hz, consistent with previous studies^23,47,48^. Although the model estimates the exponent (absolute of the spectral slope), offset and knee separately (*Methods*, eq. 1), these parameters are often strongly correlated in empirical data, given the underlying rotation of the power spectrum and the mathematical dependence between knee and slope (*Methods*, eq. 2). Note that a flattening of the power spectrum corresponds to a slope increase (values closer to 0), while a steepening of the power spectrum reflects a slope decrease (more negative values). Here, the spectral slope was tightly correlated with both offset (rho = -0.97, p < 0.0001, Pearson correlation) and knee frequency (rho = -0.73, p < 0.0001), indicating substantial redundancy. We therefore focused on the spectral slope, in line with common practices^36,37,41,43,46^. Locomotion increased pyramidal cell spiking at the pseudo-population level relative to quiescence (**Fig. 2a**; p < 0.0001, z = 12.68; Wilcoxon signed rank test), an effect corroborated by a linear mixed effects model (LME; p < 0.0001, t = - 19.03) that considered the contribution of individual mice and neurons. A similar increase was observed for (putative) fast-spiking interneurons (**Fig. 2b**; p < 0.0001, z = 9.14; Wilcoxon signed rank test) and confirmed with an LME (p < 0.0001, t = -17.87).

**Figure 2.**
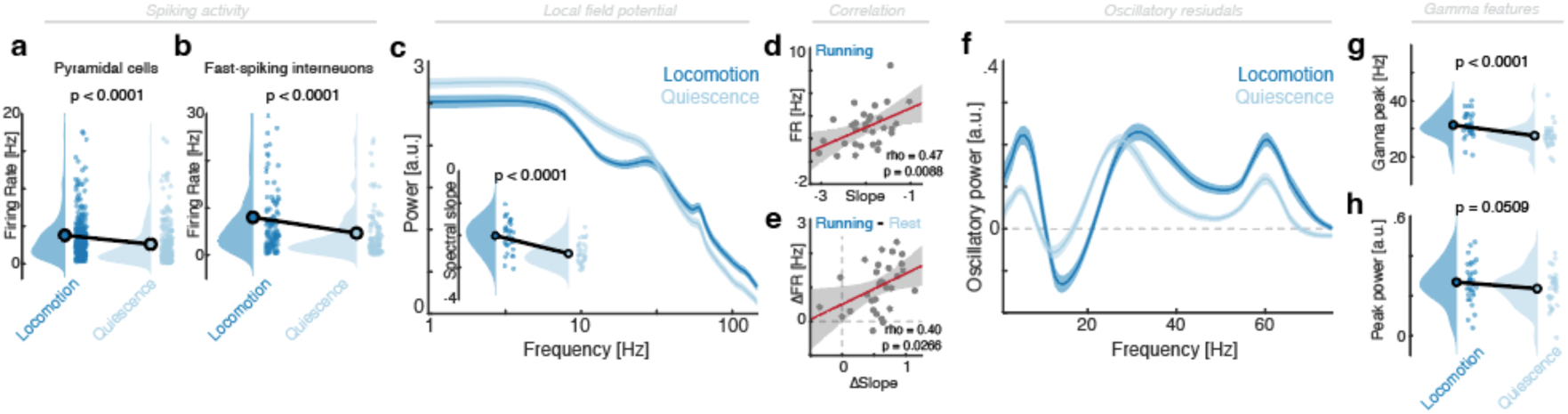
State-dependent modulation of cortical dynamics. (**a**) Statistically significant increase in putative pyramidal cell spiking during locomotion compared to quiescence. (**b**) Putative fast spiking interneurons increase their spiking during locomotion (same conventions as in panel **a**). (**c**) Power spectra for locomotion (dark blue) and quiescence (light blue). Note the overall tilt of the power spectrum with decreased low frequency power and increased high frequency power. Inset (lower left): This tilt was the result of a statistically significant change of the spectral slope. (**d**) Linear correlation between the absolute spectral slope and the pyramidal cell spiking rate. (**e**) Linear correlation between the slope and spiking rate difference. (**f**) Oscillatory residuals (full spectrum *minus* aperiodic fit; cf. panel **c**). Note the three distinct peaks in the theta (∼5 Hz), visual (∼35 Hz) and fast gamma (∼60 Hz) range. (**g**) Locomotion significantly increased the visual gamma peak frequency, (**h**) but not its power.

In parallel, the LFP power spectra exhibited a pronounced ‘tilt’, indexed by a flattening of the power spectrum (slope increase) during locomotion (**Fig. 2c**; p < 0.0001, t32 = 11.30, d = 1.97; paired t-test; *locomotion*: -2.02 ± 0.08; *quiescence*: -2.57 ± 0.06; mean ± SEM; also evident without visual stimulation, cf. **Fig. S2**). The spectral slope directly correlated with pyramidal cell spiking during locomotion (**Fig. 2d**, rho = 0.47, p = 0.0088; Pearson correlation) and quiescence (rho = 0.40, p = 0.0306), and the locomotion-induced slope change (locomotion *minus* quiescence) tracked the corresponding change in spiking between both cortical states (**Fig. 2e**; rho = 0.40, p = 0.0266).

In the oscillatory residuals, distinct peaks were apparent in the theta (< 10 Hz), visual gamma (25-40 Hz), and locomotion-associated fast gamma (∼60 Hz) range (**Fig. 2f** and **Supplementary Text**). Note, the simultaneous presence of low and fast gamma rhythm resulted from averaging across visual stimulation conditions (see below, cf. **Fig. 3/4**). Locomotion increased visual gamma peak frequency (**Fig. 2g**; p < 0.0001, t32 = 5.18, d = 0.90; *locomotion*: 31.28 ± 0.76 Hz; *quiescence*: 27.46 ± 0.70 Hz; mean ± SEM), while gamma power was not statistically significantly modulated (**Fig. 2h**; p = 0.0509, t32 = 2.03, d = 0.35; *locomotion*: 0.27 ± 0.02, *quiescence*: 0.24 ± 0.02; mean ± SEM). Together, this set of findings demonstrates locomotion-related changes in neural spiking, aperiodic activity, and oscillatory dynamics, and indicates a positive association between pyramidal cell spiking and aperiodic slope. We therefore prioritized locomotion for subsequent analyses to maximize usable data, as not all mice remained quiescent for sufficient durations.

**Figure 3.**
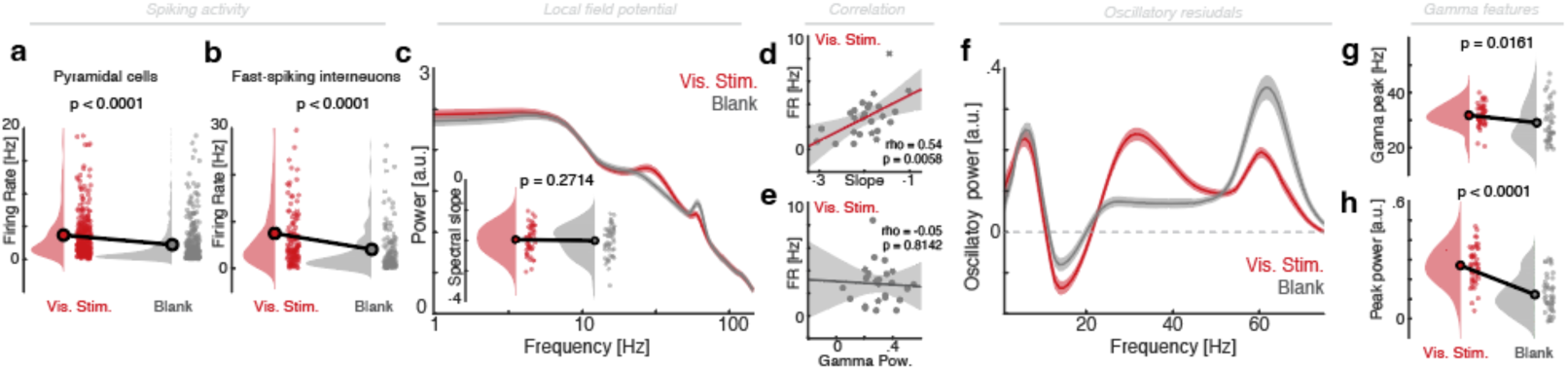
Context-dependent modulation of cortical dynamics. (**a**) Statistically significant increase in putative pyramidal cell spiking during visual stimulation (gratings) compared to no stimulation (blank screen). (**b**) Putative fast spiking interneuron spiking increases during stimulation (same conventions as in panel **a**). (**c**) Power spectra for visual stimulation (dark red) and no stimulation (grey). Note the selective modulation of the oscillatory peaks rising above the 1/f decay function. Inset (lower left): Conversely, the spectral slope did not differ significantly between the two conditions. (**d**) Significant linear correlation between the absolute spectral slope and the pyramidal cell spiking rate. (**e**) The visual gamma power (cf. panel **h**) was not significantly correlated with pyramidal cell spiking. (**f**) Oscillatory residuals (full spectrum *minus* aperiodic fit; cf. panel **c**). Note the modulation in the visual (∼35 Hz) gamma range. (**g**) Visual input significantly increased the visual gamma peak frequency and (**h**) its peak power.

To further disentangle their precise relationships, we next compared neural spiking, aperiodic and oscillatory activity during visual stimulation to a baseline condition without visual stimulation (isoluminant grey: blank screen). Visual input increased both pyramidal cell spiking (**Fig. 3a**; p < 0.0001, z = 10; Wilcoxon signed rank test) and fast-spiking interneuron spiking (**Fig. 3b**; p < 0.0001, z = 6.71), confirmed with LMEs (pyramidal cells: p < 0.0001, t = -14.19; interneurons: p < 0.0001, t = -9.44). Unlike the robust state-dependent spectral tilt (cf. **Fig. 2c**), visual stimulation did not significantly modulate the aperiodic slope (**Fig. 3c**; p = 0.2714, t47 = 1.11, d = 0.16; paired t-test; *visual stimulation*: -1.97 ± 0.07, *no stimulation*: -2.01 ± 0.07; mean ± SEM). Nevertheless, across animals, the individual spectral slope significantly correlated with pyramidal spiking rates during visual stimulation (**Fig. 3d**; rho = 0.54, p = 0.0058, Pearson correlation) and also without stimulation (rho = 0.51, p = 0.0094), whereas visual gamma power did not (**Fig. 3e**; rho = -0.05, p = 0.8142; defined on oscillatory residuals, cf. **Fig. 3f**). Clear visual gamma peaks emerged only during visual stimulation (**Fig. 3f**), with modestly higher peak frequency (**Fig. 3g**; p = 0.0161, t47 = 2.50, d = 0.36; *visual stimulation*: 31.78 ± 0.57 Hz; *no stimulation*: 29.07 ± 1.02 Hz; mean ± SEM) and markedly elevated peak power (**Fig. 3h**; p < 0.0001, t47 = 10.85, d = 1.57; *visual stimulation*: 0.27 ± 0.01; *no stimulation*: 0.12 ± 0.01; mean ± SEM).

We next varied stimulus size during visual stimulation. As expected, pyramidal (**Fig. 4a**, p < 0.0001, χ^2^ = 265.65; Friedman test) and interneuron (**Fig. 4b**; p < 0.0001, χ^2^ = 185.09) spiking showed a non-monotonic dependence on size, where neural spiking is increased for stimuli that match the receptive field size (*second* column in **Fig. 4a/b**), but then reduced for larger stimuli, consistent with surround suppression. At the LFP level (**Fig. 4c**), oscillatory dynamics were strongly modulated, whereas the aperiodic spectral slope was not (**Fig. 4d**; p = 0.4369, F3,135 = 0.91; GG-RM-ANOVA). Oscillatory peak frequency and power scaled linearly with increasing stimulus size (**Fig. 4e**). Visual gamma peak frequency decreased with larger stimuli (**Fig. 4f**; p < 0.0001, F3,148 = 9.86; from 36.0 to 30.6 Hz), while gamma power increased (**Fig. 4g**; p < 0.0001, F2,114 = 108.32; from 0.11 to 0.43). This establishes a triple dissociation between neural spiking, aperiodic and oscillatory dynamics: spiking varies non-monotonically with stimulus size, aperiodic dynamics remain stable, and oscillatory gamma activity scales with size. Subsequent analyses therefore focused on the largest visual stimulus condition to maximize oscillatory gamma amplitude (cf. **Fig. 4e**).

**Figure 4.**
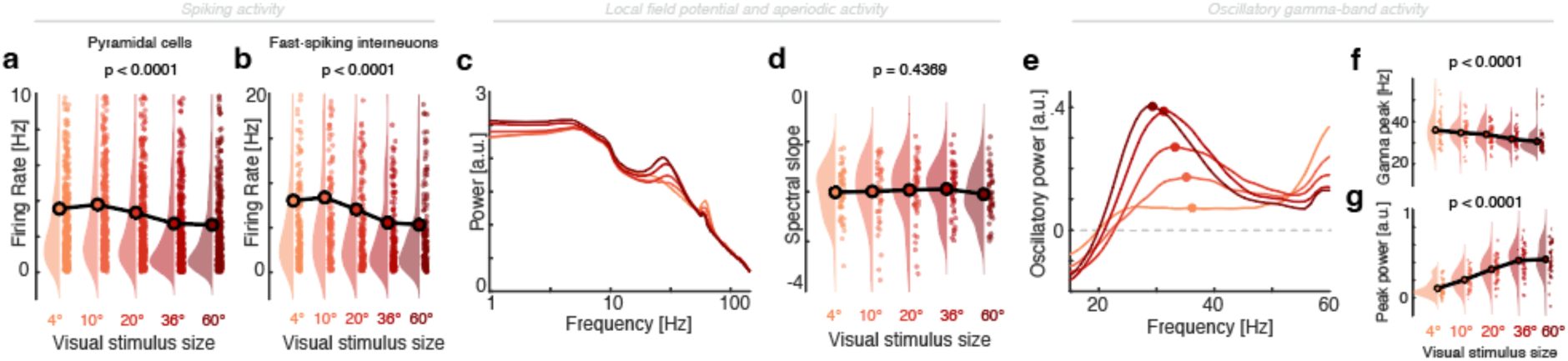
Stimulus-dependent modulation of cortical dynamics. (**a**) Statistically significant non-monotonic modulation of putative pyramidal cell spiking relative to different stimulus sizes (p < 0.0001; increasing size from left to right). Note the strongest response (second bin) for stimuli that matched the receptive field size. (**b**) Putative fast spiking interneurons follow the same pattern as pyramidal cells (p < 0.0001; same conventions as in panel **a**). (**c**) Power spectra for the five different visual stimulation conditions (same conventions as in panel **a**). Note that the modulation was primarily confined to the oscillatory peaks rising above the 1/f decay function. (**d**) Conversely, the spectral slope did not differ significantly between the five stimuli (p = 0.4369). (**e**) Oscillatory residuals (full spectrum *minus* aperiodic fit; cf. panel **c**). Note the modulation in the visual gamma response (colored points indicate the average peak). (**f**) Increasing stimulus size significantly reduced the visual gamma peak frequency (p < 0.0001) and (**g**) significantly increased its amplitude (p < 0.0001).

To causally probe the relationship between neural spiking and LFP dynamics, we optogenetically suppressed interneuron subtypes during visual stimulation. Suppressing **SST**-positive interneurons (a direct monosynaptic inhibitor of pyramidal cell dendrites; cf. **Fig. 1a**) reduced overall inhibition and thereby increased pyramidal cell spiking (**Fig. 5a**; p < 0.0001, z = -6.5; Wilcoxon signed rank test; confirmed with an LME: p < 0.0001, t = 6.79) and interneuron activity (**Fig. 5b**, p = 0.0002, z = -3.77; Wilcoxon signed rank test; confirmed with an LME: p < 0.0001, t = 4.12). This was accompanied by flattening of the LFP power spectra (**Fig. 5c**; cf. **Fig. S3**), which corresponded to an increase of the spectral slope (**Fig 5d**; p = 0.0009, t16 = -4.07, d = -0.99; *light off*: -2.02 ± 0.13, *light: on*: - 1.48 ± 0.14; mean ± SEM). In oscillatory residuals (**Fig. 5e**; cf. panel **c**), visual gamma-band peak frequency was not significantly altered (**Fig. 5f** *upper panel*; p = 0.0514, t16 = -2.11, d = -0.51; *light off*: 30.10 ± 0.71 Hz; *light on*: 31.82 ± 1.01 Hz; mean ± SEM), but gamma power was strongly reduced (**Fig. 5f** *lower panel*; p < 0.0001, t16 = 6.58, d = 1.60; *light off*: 0.45 ± 0.04, *light on*: 0.31 ± 0.03), as demonstrated previously^23^.

**Figure 5.**
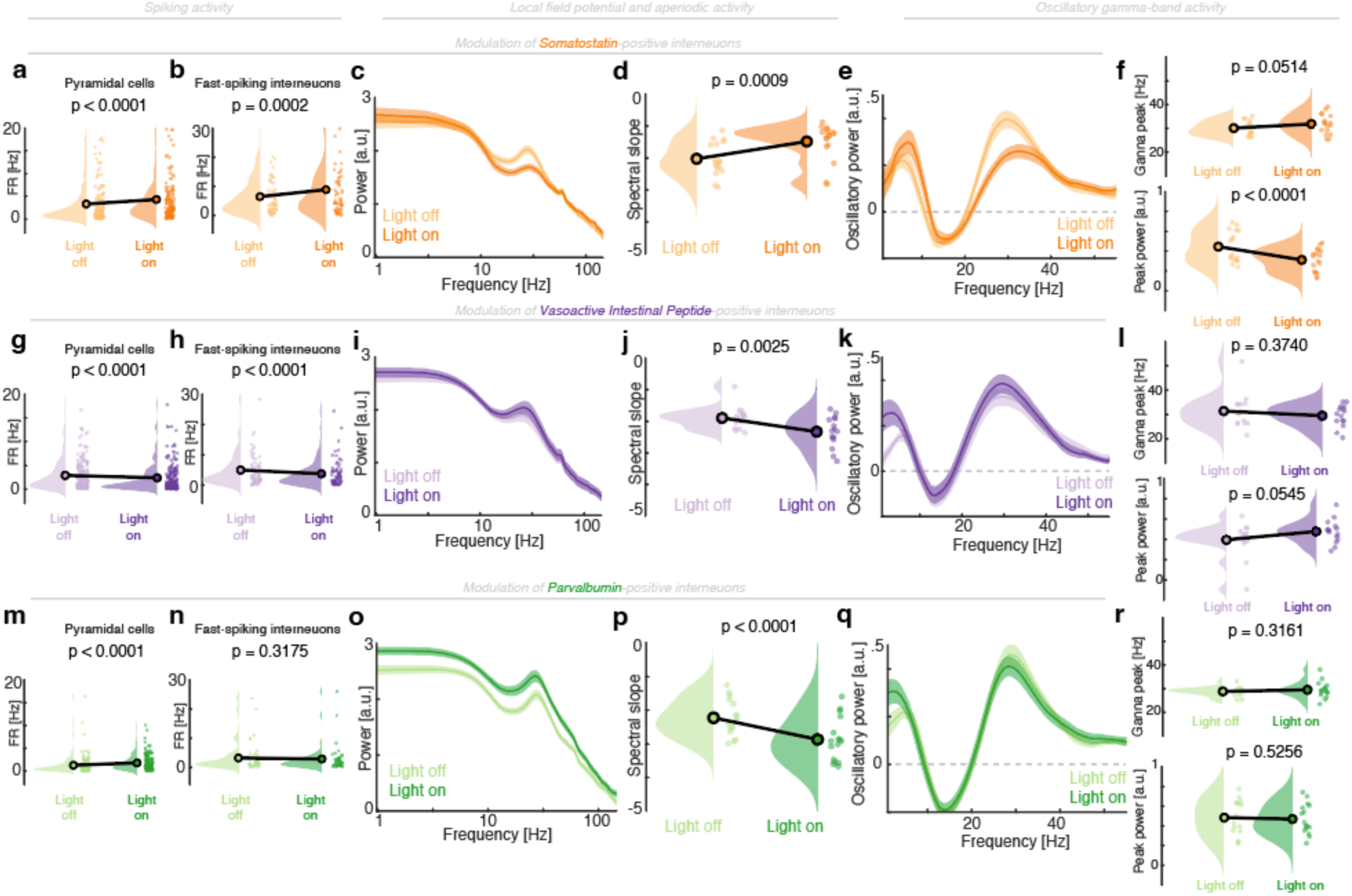
Optogenetic suppression of interneurons modulates aperiodic and oscillatory features. (**a-f**) Inactivation of SST-positive interneurons. (**a**) Statistically significant increase in putative pyramidal cell spiking during optogenetic suppression (dark orange) as compared to the control condition (light orange). (**b**). Fast-spiking interneurons. (**c**) Power spectrum during light off and on. (**d**) Statistically significant modulation of aperiodic activity (spectral slope), which corresponds to a flattening of the power spectrum (cf. panel **c**). (**e**) Oscillatory residuals. (**f**) Visual gamma peak frequency (top). A significant modulation of gamma peak amplitude (bottom) was observed. (**g-l**) Suppression of VIP-positive interneurons. Same conventions as in panels **a-f**. Note the significant decrease of neural spiking (panel **g/h**) as well as a significant steepening of the spectral slope (panel **j**). (**m-r**) Suppression of PV-positive interneurons. Same conventions as in panels **a-f**. Note the increase in pyramidal cell spiking (panel **m**) with a concomitant decrease in the spectral slope (panel **p**).

Next, we suppressed **VIP**-positive interneurons, which mainly target SST-positive interneurons (**Fig. 1a**) and thus inhibit inhibitory drive, which led to a net increase in excitation. Suppressing VIP-positive interneurons disinhibited SST interneurons that then decreased pyramidal (**Fig. 5g**; Wilcoxon signed rank test: p < 0.0001, z = 7.63; LME: p < 0.0001, t = -8.75) and fast-spiking interneuron spiking (**Fig. 5h**; signed rank test: p < 0.0001, z = 5.21; LME: p < 0.0001, t = -5.84). At the LFP level, this excitation decrease translated to a steepening of the spectrum; i.e., a reduction of the spectral slope (**Fig. 5i/j**; p = 0.0025, t14 = 3.68, d = 0.94; *light off*: -1.90 ± 0.09, *light on*: -2.33 ± 0.13; mean ± SEM), but left oscillatory dynamics largely unchanged (**Fig. 5k/l**; gamma peak frequency: p = 0.3740, t14 = 0.92, d = 0.24; *light off*: 31.38 ± 1.81 Hz, *light on*: 29.49 ± 1.14 Hz). We only observed a discrete, yet statistically non-significant increase in gamma-band power (**Fig. 5l** *lower panel*; p = 0.0545, t14 = -2.10, d = -0.54; *light off*: 0.40 ± 0.04, *light on*: 0.48 ± 0.03).

Lastly, we also inferred the contribution of **PV**-positive interneurons, which mainly target the pyramidal cell soma (**Fig. 1a**). Suppressing PV inhibitory interneurons increased pyramidal cell spiking (**Fig. 5m**; signed rank test: p < 0.0001, z = -9.34; LME: p < 0.0001, t = 10.32), but not fast-spiking interneuron activity (**Fig. 5n**, signed rank test: p = 0.3175, z = -1; LME: p = 0.8231, t = 0.22). In the LFP spectral domain (**Fig. 5o**), suppression of PV interneurons produced a surprisingly strong opposite tilt and steepening of the power spectrum; thus, decreasing the spectral slope (**Fig. 5p**; p < 0.0001, t15 = 6.84, d = 1.71; *light off*: -2.23 ± 0.13, *light on*: -2.87 ± 0.16) without significant changes in oscillatory features (**Fig. 5q**), such as gamma-band peak frequency (**Fig. 5r**; p = 0.3161, t15 = -1.04, d = -0.26; *light off*: 28.81 ± 0.47 Hz, *light on*: 29.48 ± 0.86 Hz) or peak power (p = 0.5256, t15 = 0.65, d = 0.16; *light off*: 0.48 ± 0.04, *light on*: 0.47 ± 0.04). Interestingly, this set of results contradicted the notion that pyramidal cell spiking is always modulated in the same direction as the spectral slope (as implied in e.g., **Fig. 5a/d** or **Fig. 5g/j**). While a relationship between spiking and the spectral slope had been observed during locomotion (**Fig. 2a/c**), the suppression of PV-positive interneurons dissociated spiking from slope, demonstrating that pyramidal cell spiking does not invariably covary with the aperiodic slope.

Finally, we directly quantified the three-way relationship between neural spiking, aperiodic slope and gamma synchrony. At baseline (*light off*), the spectral slope was only weakly correlated with pyramidal cell spiking across all available mice (**Fig. 6a**, rho = 0.22, p = 0.1384, Spearman correlation). In contrast, optogenetically-induced changes of cell subtype activity cleanly separated experimental conditions (light on *minus* light off; **Fig. 6b**) and revealed a significant association of spectral slope modulation with pyramidal cell spiking modulation at the group level (rho = 0.41, p = 0.0045, Spearman correlation). Modulation of SST- and VIP-positive interneurons lay *on*-diagonal (consistent with comodulation), whereas suppression of PV-positive interneurons gave rise to a prominent *off*-diagonal distribution (upper left quadrant in **Fig. 6b**), again, highlighting that increased neural spiking was accompanied by a steeper spectrum and a decrease of spectral slope (cf. **Fig. 5m/p**). We hypothesized that this divergence reflected changes in neural synchrony in addition to changes in overall excitability. Therefore, we assessed neural synchrony by means of visual gamma-band power during optogenetic suppression. Indeed, baseline gamma-band power did not correlate with neural spiking (**Fig. 6c**; rho = -0.05, p = 0.7580, Spearman correlation), but optogenetically induced changes (light on *minus* light off) revealed a robust inverse relationship: larger decreases in gamma power (less synchrony) predicted larger increases in pyramidal cell spiking (**Fig. 6d**; rho = -0.48, p = 0.0009, Spearman correlation). Together, these observations indicate that neural spiking is modulated by both aperiodic and oscillatory gamma activity. Qualitatively and quantitatively highly similar results were observed when we averaged across all visual stimulation conditions (**Fig. S4**).

**Figure 6.**
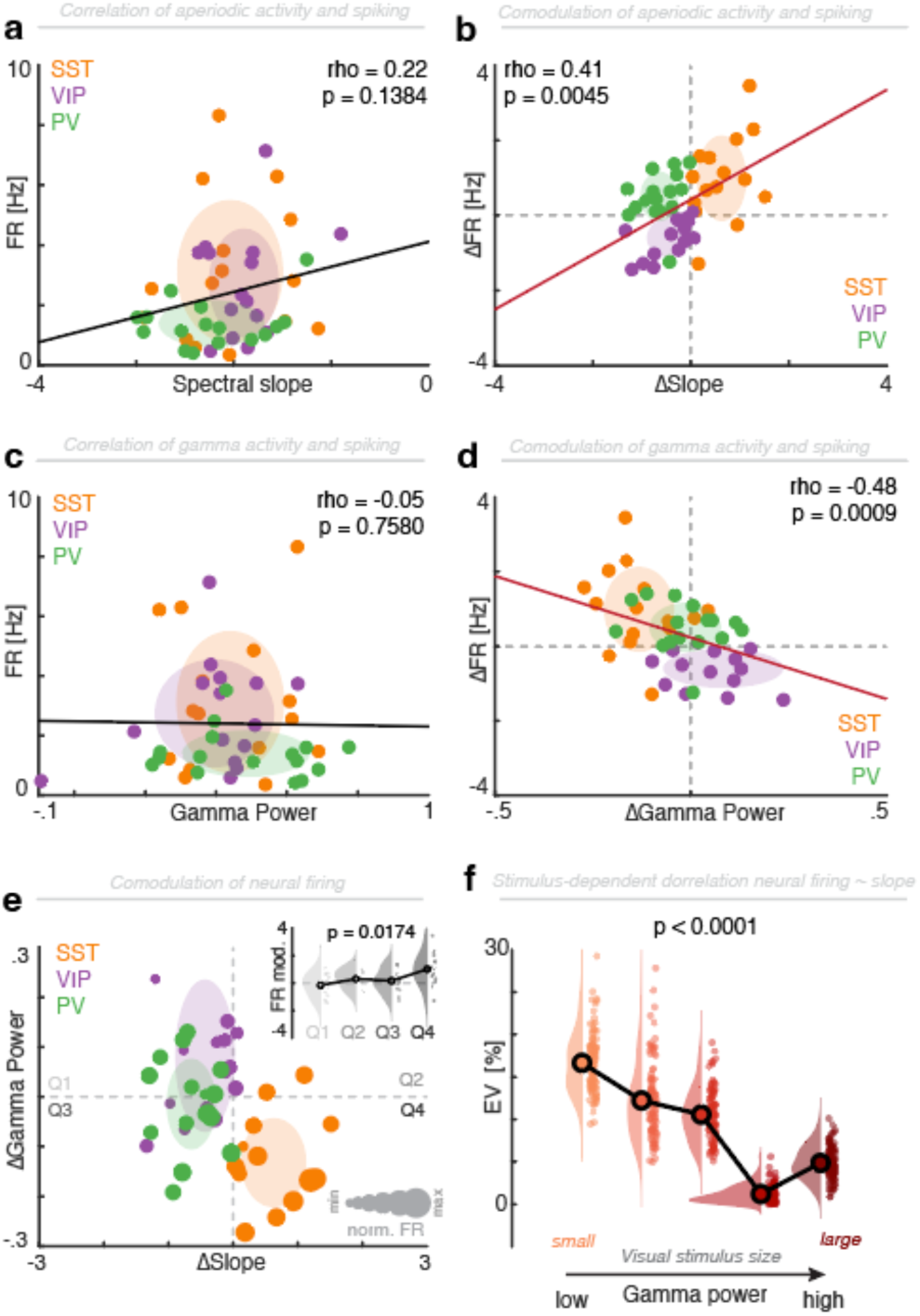
Modulation of interneuron activity shapes aperiodic and oscillatory activity. (**a**) Weak correlation between the spectral slope and pyramidal cell spiking. Flatter spectral slopes signal increased neural spiking. (**b**) Optogenetic modulation (light on *minus* light off) demonstrates a significant relationship with a clear differentiation and compartmentalization of the different interneuron cell types. While suppression of SST-positive interneuron led to more spiking and flatter spectral slopes (upper right quadrant), suppressing VIP-positive interneurons gave rise to the opposite pattern (lower left quadrant) with decreased spiking rates and steeper spectral slopes. A dissociation between spiking and the spectral slope was observed for PV-interneuron suppression (upper left quadrant), where an increase in spiking was accompanied by a steepening of the power spectrum. (**c**) Visual gamma power was not correlated with the overall spiking rate. (**d**) However, the optogenetically-induced gamma power modulation (light on *minus* light off) demonstrated a significant negative correlation, highlighting that increases in visual gamma synchrony predicted spiking rate decreases. (**e**) The paradoxical result for suppression of PV-positive interneurons (cf. panel **b**) can be attributed to a simultaneous modulation of aperiodic (x-axis) and oscillatory (y-axis) activity. The size of the colored circles corresponds to relative spiking rate, with increases for SST- (orange) and PV-positive (green) interneurons and decreases for VIP-positive interneurons (purple). Inset (Upper right): Statistical comparison of the spiking rate modulation in the four quadrants, highlighting that the strongest increase in spiking was observed when the spectral slope increased (flattening of the power spectrum), while gamma synchrony decreased (Q4). (**f**) The correlation (in percent explained variance, squared correlation coefficients, cf. **Fig. S5**) between neural spiking and the spectral slope (cf. panel **a**) as a function of different visual stimulation conditions that induced variable gamma-band power (cf. Fig. 4e**/g**). The relationship between aperiodic activity and neural spiking is strongly attenuated in states of strong oscillatory synchrony.

Joint visualization of normalized spiking rate modulation as a function of spectral slope and gamma-band modulation (**Fig. 6e**) revealed that especially the suppression of PV interneurons gave rise to a variable gamma-band modulation. For statistical quantification (*inset* **Fig. 6e**), we grouped the spiking rate modulation into four quadrants and observed that the strongest increases in neural spiking occurred when the spectral slope increased (flatter power spectrum) while gamma-band synchrony decreased (*lower right Quadrant 4;* p = 0.0174, F3,42 = 3.77, ANOVA).

Finally, during baseline activity, the relationship of neural spiking and aperiodic activity weakened systematically as gamma power increased (**Fig. 6f**; 100 bootstraps, random subsampling of 90% of the subjects), dropping from rho = 0.41 to rho = 0.10 (cf. **Fig. 4e/g** and **Fig. S5**; Spearman correlation; GG-RM-ANOVA: p < 0.0001, F3.1, 310.5 = 471.7). A similar pattern was evident during optogenetic stimulation (**Fig. S5**).

In sum, these results demonstrate that aperiodic activity explains more variance in pyramidal cell spiking than gamma-band oscillations, but this effect is pronounced in states of low oscillatory synchrony (**Fig. S6**). In turn, strong gamma synchronization substantially attenuates the coupling between aperiodic activity and neural spiking. Thus, the results uncover an intricate relationship between neural spiking and oscillatory and aperiodic LFP features.

## Discussion

Here we demonstrate that optogenetic suppression of SST, VIP, and PV interneurons in mouse V1 modulates both neural spiking activity and LFP spectra, broadly consistent with predictions from simplified circuit models that link LFP features to neural excitability^36^. Although this interneuron circuit motif has primarily been used to account for gamma oscillations^23,45^, our results now demonstrate that it also explains aperiodic background activity. Across cortical states (**Fig. 2d/e**) and behavioral contexts (**Fig. 3d**), pyramidal spiking covaried with the 1/f slope of the electrophysiological power spectrum, but not with evoked gamma oscillations (**Fig. 3e**). Visual stimulation revealed a dissociation between these three signatures: stimulus-size changes produced non-monotonic modulation of neural spiking (**Fig. 4a/b**) alongside systematic modulation of gamma synchrony (**Fig. 4f/g**), while leaving aperiodic activity largely unchanged (**Fig. 4d**).

Causal circuit perturbations further refined this picture. Optogenetic suppression of dendrite-targeting SST interneurons increased spiking and flattened the power spectrum (increased the spectral slope), whereas suppressing VIP interneurons produced the opposite pattern (**Fig. 5**), aligning with model-based predictions for increased versus decreased cortical excitability. In contrast, suppressing soma-targeting PV interneurons dissociated spiking from aperiodic activity (**Fig. 5m/p**). This apparent discrepancy is explained by a joint dependence of spiking on both oscillatory synchrony and aperiodic activity: the largest increases in spiking occurred when the 1/f slope increased while gamma synchrony decreased (**Fig. 6e/f**). More generally, aperiodic activity was most predictive of spiking rate in conditions with low oscillatory synchrony, underscoring that the relationship between aperiodic activity and spiking, and by extension its interpretation as a proxy for neural excitability, depends strongly on behavioral context and network state.

### The cellular correlates of extracellular fields in the gamma range

Gamma oscillations are frequently implicated in information coding and interareal communication^3,12,13,26,28^, with much of the mechanistic evidence stemming from animal models^9,12,13^. In human neurophysiology, however, it has long been recognized that broadband power increases in the gamma range must not be conflated with stimulus-specific narrow-band gamma oscillations^25,28^. Fueled by novel tools for spectral parametrization^46^, it has become evident that apparent increases of power in the gamma frequency range might be largely driven by a flattening of the power spectrum (spectral slope increase) rather than enhanced rhythmic synchrony. Consistent with this, our results demonstrate that broadband power increases largely index changes in the 1/f slope (**Fig. 2**), in line with cellular-resolution evidence from two-photon imaging linking pyramidal cell activity to aperiodic dynamics^43^. By contrast, visual stimulation with gratings predominantly modulated narrow-band gamma oscillations (**Fig. 3**) without a measurable change in 1/f slope. Strikingly, neural spiking still correlated more closely with the aperiodic 1/f slope than with gamma synchrony (**Fig. 3d/e**). Systematically varying visual stimulation conditions further dissociated neural spiking, aperiodic activity and oscillations (**Fig. 4**), demonstrating that LFP spectra cannot be explained by a simple linear superposition of neural spiking patterns. Finally, by optogenetic suppression of interneuron subtypes, we provide causal evidence that distinct inhibitory pathways differentially shapes LFP spectra (**Fig. 5**): SST suppression increased neural spiking and flattened the power spectrum (increased the spectral slope; **Fig. 5a/d**), whereas VIP suppression decreased spiking and steepened the power spectrum (decreased the 1/f slope; **Fig. 5g/j**). These two findings directly confirm several empirical observations in human intracranial and scalp EEG linking flatter power spectra to increased excitability^36,41,43,49–51^, and vice versa, especially given that LFP and EEG signals largely reflect postsynaptic dendritic potentials^1,5^.

The interneuron circuit motif (**Fig. 1a**) was initially proposed based on computational modeling of prefrontal cortex dynamics^44^ and later repeatedly linked to the generation of gamma oscillations^23,24^. Here, we extend its relevance to aperiodic activity. It has also long been recognized that the precise composition of the motif on the population level (cf. **Fig. 1c**) might change along the cortical hierarchy as the ratio of SST to PV cells increases when moving from primary sensorimotor to association areas^45,52^. This change along the cortical hierarchy might be mirrored in the hierarchical gradient of 1/f aperiodic activity in the human brain^46,53^. However, the divergent effects of SST and PV suppression on the neural spiking and spectral slope relationship (**Fig. 5**) warrant caution when interpreting aperiodic activity, indicating that aperiodic metrics can be shaped by multiple inhibitory mechanisms, not a single excitability axis.

### The neurophysiologic basis of aperiodic activity

Aperiodic (broadband) activity has periodically been appreciated over the years to complement, falsify or expand oscillation-centric views^30,36,54,55^, and interest has accelerated with recent empirical observations, novel computational models^35,36^, analytic approaches^46^, and theoretical frameworks^49,56^. Yet the cellular basis of aperiodic activity remains unknown, as most studies rely on macroscale data that indirectly infer the link to neural excitability dynamics^37,41^. Pharmacological, in vivo imaging, and invasive electrophysiology studies jointly support a link between aperiodic activity and neural excitability, but not necessarily a direct readout of the balance of excitation and inhibition as suggested initially^39,42,43,50,57^. This apparent discrepancy that may reflect model assumptions such as the omission of recurrent connections between excitatory and inhibitory neurons^36^. In our recordings, spiking of putative pyramidal and fast-spiking interneurons (which presumably mainly reflected PV interneurons) covaried across state and stimulus conditions (cf. **Fig. 2a/b**, **Fig. 3a/b** and **Fig. 4a/b**). In sum, our results provide a causal, cell-type-informed basis for interpreting aperiodic activity, firmly grounded on predictions derived from a canonical microcircuit motif, and thereby open new avenues to relate broadband spectral features to circuit excitability across behavioral contexts, vigilance, and disease states.

### Gamma oscillations shape the coupling between aperiodic activity and spiking

Spectral parametrization separates oscillatory from aperiodic components of the LFP spectrum (**Fig. 1e**), implicitly suggesting partially dissociable neural mechanisms. Our results reveal an intricate interplay: The correlation between neural spiking and spectral slope was strongest when oscillatory gamma activity was low and was markedly attenuated as gamma synchrony increased (**Fig. 6e/f**), suggesting that synchrony can decouple spike rate from broadband spectral features, possibly by constraining spike timing. Mechanistically, gamma oscillations are typically viewed as an emergent property of local excitation-inhibition interactions^9,12^ (e.g., PING - pyramidal-interneuron gamma), in which recurrent excitation recruits fast inhibition to produce rhythmic cycles of pyramidal spiking and interneuron-mediated suppression. While PV+ interneurons are widely considered central to gamma generation^16,21^, recent work also implicates SST+ and VIP+ interneurons through their influence on dendritic integration and disinhibitory gating^23,24^. Here, we reproduce classic state- and stimulus-dependencies of gamma, including locomotion, arousal, task engagement, and stimulus drive (**Fig. 2-4**), consistent with context-dependent shifts in population excitation/inhibition (E/I-) balance. The results are also in accordance with the idea that SST+ contribute to gamma synchrony in V1^23^ (cf. **Fig. 5f/l/r**). We primarily focused on pyramidal cell spiking as the number of recorded interneurons precluded a reliable estimation of the balance of excitation and inhibition. As excitability and E/I-balance are central to both gamma oscillations and aperiodic activity, future studies need to clarify how strongly these signatures overlap mechanistically and how they dynamically interact to support context-dependent, goal-directed circuit operations underlying complex behaviors and cognition.

### Limitations

This study uses a between-group design across 60 mice recorded exclusively in V1, so extension to other cortical regions and within-subject designs will be important. We did not assess inter-areal gamma coherence, which might constitute a dissociable phenomenon from local gamma power. While optogenetic inactivation targeted specific cell types, our electrophysiological recordings only enabled separation into putative pyramidal and fast-spiking interneurons based on the spiking and waveform characteristics. Interpreting optogenetic suppression of PV+ interneurons is also nontrivial. Because PV+ cells stabilize recurrent networks by regulating pyramidal-cell population firing rates, suppression was intentionally kept modest to avoid network destabilization and seizure-like activity. Under these conditions, incomplete silencing and recruitment of non-opsin-expressing (or weakly expressing) PV+ neurons may provide compensatory inhibition, potentially accounting for the relatively small changes observed in the fast-spiking population. Lastly, this study was conducted in rodents, thus cross-species translations to humans should be made cautiously, even though large-scale gradients appear conserved ^58–60^.

## Conclusions

Collectively, our results link aperiodic LFP activity to identifiable microcircuit substrates and show how state-dependent excitability changes shape both the aperiodic background activity and gamma oscillations. By integrating cell-type specific perturbations with spectral parametrization and spiking measurements, we provide a biologically grounded, mechanistic framework for interpreting LFP spectra in terms of synaptic and circuit dynamics across behavioral contexts. This framework offers more specific inferences from population signals and strengthens the translational use of aperiodic activity as a marker of altered neural excitability across a range of conditions^61^, including disorders-of-consciousness^41,62,63^, epilepsy^64,65^, and movement disorders^51,66^.

## Material and Methods

Here we reanalyzed neural spiking activity and local field potentials from two previously reported studies. The full methodological details of data acquisition and preprocessing are reported in^23,24^ and summarized below. In brief, we pooled the available source data into a dataset that consisted of 60 mice (n = 20 SOM-Cre, n = 21 VIP-Cre and n = 19 PV-Cre) who underwent the same experimental protocol, which included five visual stimulation conditions (gratings of different sizes) as well as one condition without visual input (isoluminant gray screen) during two different contexts (locomotion/quiescence), while specific interneurons were either optogenetically inactivated or not. For all animals, we obtained grand-average power spectra from the local field potential for all conditions [6 (visual conditions) x 2 (states) x 2 (light on/off)] as well as spiking activity (N = 832 cells) from putative pyramidal cells (N = 610) as well as from fast-spiking interneurons (N = 205), which were used for all analyses. 17 cells had intermediate spike widths and could thus not be further differentiated. Note that not all animals yielded usable data for all conditions, as in some instances, too few trials were collected in either the locomotion or quiescent state, no responsive neurons were identified or the grand-average power spectra exhibited residual noise. Hence, we sub-selected data for different analyses based on availability.

The full experimental procedures are reported elsewhere. In brief, animals were neonatally injected with either eNpHR3.0-YFP or AAV-EF1a-DIO-eArch3.0-YFP. After expression of the virus and habituation to the running wheel, a small craniotomy was opened over V1 and animals were recorded with 16-channel silicon probes (NeuroNexus, A1×16-5mm-25-177-A16). Visual stimuli consisted of drifting square-wave gratings at 0.04 cycles per degree and 2 cycles per second centered on the average MUA receptive field presented for 2s with at least 1s inter-stimulus interval. Gratings were presented in different configurations, here we focused on full contrast gratings of eight different directions (0°–315° in steps of 45°) and five different sizes (4, 10, 20, 36, and, if possible, 60 visual degrees; if the RF was not perfectly centered on the monitor, the effective largest size was slightly smaller). In addition, we included a condition without visual stimulation, where the screen showed an isoluminant (to the gratings) gray. Gratings drifted for 2s with at least 1s inter-trial intervals with the optogenetic LED switched on for 1 s starting 0.5 s after the start of the visual stimulus in 50% of the trials. The period of light was chosen to influence the stable steady-state of the response to the grating and all analysis was performed during this time window. Local field potentials were extracted by low-pass filtering the raw signal, sampled at 30 kHz, below 200 Hz and subsequent down-sampling to 1 kHz. We analyzed the LFP from the electrode contact closest to a cortical depth of ∼350 mm (in cortical layer 3). The power spectra were computed in an 800 ms analysis window starting 200 ms after light onset (to exclude any photo-electric artifacts sometimes present in the first ∼150 ms after light onset) using multi-taper estimation as implemented in the Chronux package (http://chronux.org/) with three tapers. All resulting grand-average spectra were visually inspected. Spectra that exhibited noise were manually flagged and removed from subsequent analyses.

### Spectral parametrization

We parametrized all spectra and obtained the aperiodic component using the Specparam/FOOOF toolbox through the Matlab wrapper function with standard parameters ^46^. We parametrized all spectra with the ‘knee’ option in the range from 1-120 Hz. All other parameters were set to default values. Aperiodic activity was defined by its slope parameter *χ*, the offset *c* and a constant *k* (reflecting the knee parameter) as

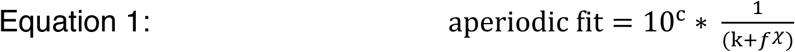

The relationship of the knee parameter *k* and the knee frequency ƒk is given by

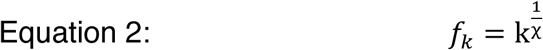

In case that the algorithm did not converge or no plausible knee parameter was identified, the parametrization was repeated using a linear fit (‘fixed’ mode, equivalent to *k* = 0 in equation 1). We only extracted the aperiodic parameters from the FOOOF output for subsequent analyses. We calculated the aperiodic background fit from the identified offset, time constant and exponent (equation 1). We multiplied the exponent with -1 to obtain the spectral slope, which we reported throughout to highlight the negative 1/f decay function of the power spectrum. The goodness-of-fit (*R*^2^) was obtained from the FOOOF model. We defined the oscillatory residuals as the difference of the original power spectrum minus the aperiodic background fit. This last step deviated from the typical FOOOF implementation as this facilitated visualization of the oscillatory component and consistent peak detection to delineate gamma-band oscillations. We detected the oscillatory gamma peak and amplitude on the oscillatory residuals using the findpeaks.m function in Matlab in the range from 15-55 Hz after smoothing the spectra with a 5 Hz sliding average function. We defined NPeaks = 1 and MinPeakWidth = 3 Hz. In case this did not yield a plausible solution, the estimation was repeated without requiring a minimum width of the identified peak. To account for the smaller gamma-band response for the smallest visual gratings, we also adjusted the frequency range (35-55 Hz) if necessary. To identify the theta peak frequency, we employed a comparable approach, but defined the frequency range as 1-15 Hz with MinPeakWidth = 1 Hz.

### Quantification and statistical analysis

We employed paired and unpaired statistics at the subject and pseudo-population level as indicated for every respective analysis. For group comparisons, we used paired or unpaired t-tests (e.g., **Fig. 2c/g/h**) as well as Wilcoxon signed rank tests if applicable (e.g., **Fig. 2a/b** and **Fig. 3a/b**). Spiking data was assessed at the pseudopopulation level and results were confirmed using linear mixed effect models with experimental condition (e.g., state or visual stimulation condition) as fixed effects and random intercepts for subjects and units nested within subjects (**Fig. 2a/b** and **Fig. 3a/b**). In addition, we employed one-way or repeated-measures analysis-of-variance (ANOVA; e.g., **Fig. 4a/b/d/f/g**), which were Greenhouse Geisser-corrected if applicable, the Friedman test for non-parametric, repeated-measures variance analyses, and linear correlations as indicated (e.g., **Fig. 2d**). Moreover, we employed cluster-based permutation tests (e.g., **Fig. S5**) to correct for multiple comparisons across frequencies after thresholding the primary test statistic (t-test) at p < 0.05 and randomly swapping condition labels 1000 times. The permutation p-value was then calculated against the surrogate distribution.

## Supplementary Information

### Supplementary Text

#### Quantification of spectral parametrization

We assessed the goodness-of-fit for various conditions and observed overall high model fits. Model fits were over 98% for all animal subgroups (SST: 98.5 ± 1.8%, VIP: 99.0 ± 0.6%, PV: 98.4 ± 0.7%). For locomotion, goodness-of-fit was above 98% (*locomotion*: 98.5 ± 1.8%, *quiescence*: 99.3 ± 1.1%, median ± SD). Likewise, parametrization during optogenetic inactivation remained above 98% and was on par with the light-off condition (light off: 98.8 ± 1.1%, light on: 98.8 ± 1.6%). Moreover, the different visual stimulation conditions were also well parametrized at >98% (from small to large: (1) 99.3 ± 1.4%, (2) 98.9 ± 1.6%, (3) 98.3 ± 1.6%, (4) 98.5 ± 1.4%, (5) 98.6 ± 1.0%). In addition, we observed that baseline conditions between mouse groups were qualitatively and quantitatively comparable (**Fig. S1**).

#### State-specific oscillatory signatures: Locomotion vs. quiescent rest

As highlighted in **Fig. 2f**, there were additional oscillatory peaks besides the visual gamma peak. Specifically, theta power (p < 0.0001, t32 = 5.33, d = 0.93; *locomotion*: 0.30 ± 0.03, *quiescence*: 0.15 ± 0.01; mean ± SEM; no change of peak frequency: p = 0.7811, t32 = 0.28, d = 0.05; *locomotion*: 5.09 ± 0.38 Hz, *quiescence*: 4.97 ± 0.29 Hz; mean ± SEM; detected in the range from 1-15 Hz) and fast gamma power (p < 0.0001, t32 = 7.92, d = 1.38; *locomotion*: 0.24 ± 0.02, *quiescence*: 0.13 ± 0.01; mean ± SEM; no change of peak frequency: p = 0.8517, t32 = -0.19, d = -0.03; *locomotion*: 60.93 ± 0.41 Hz, *quiescence*: 61.08 ± 0.73 Hz; mean ± SEM; detected in the range from 55-80 Hz) were also significantly increased during locomotion.

## Acknowledgements

This work was supported by the German Research Foundation (VE938/2-2).

## Supplementary Figures

**Supplementary Fig. S1.**
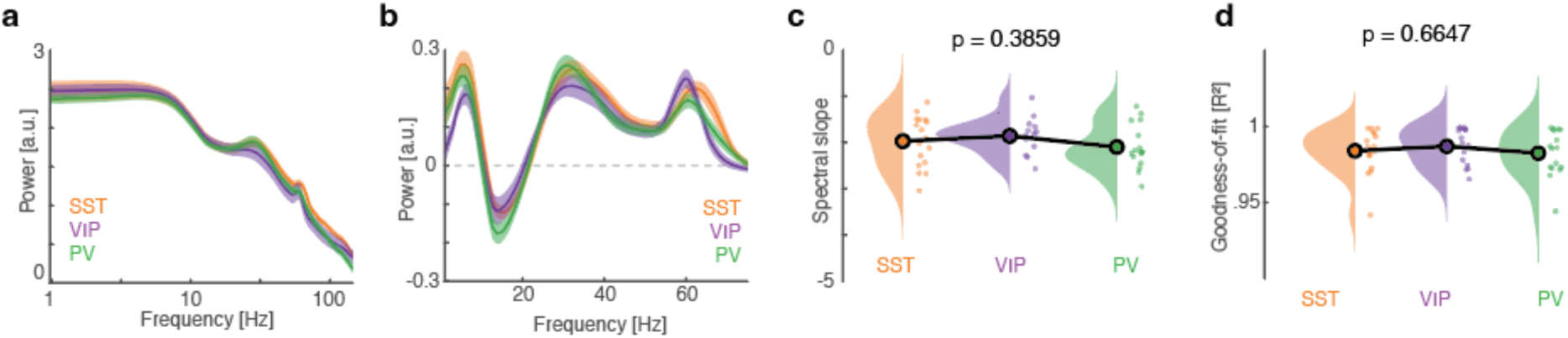
Baseline activity is comparable for SST, VIP and PV animals. (**a**) Grand-average power spectra during baseline (light off, locomotion, averaged across all stimulus conditions) did not differ between the three mouse groups (all available animals were included after artifact rejection). (**b**) Oscillatory residuals, highlighting theta, visual and fast gamma rhythms. (**c**) The spectral slope did not differ significantly between the three groups (p = 0.3859, F2,45 = 0.97, ANOVA). (**d**) Average goodness-of-fit was high (98.4 ± 1.4%) and did not differ between the three groups (p = 0.6647, F2,45 = 0.41, ANOVA). Related to **Results**.

**Supplementary Fig. S2.**
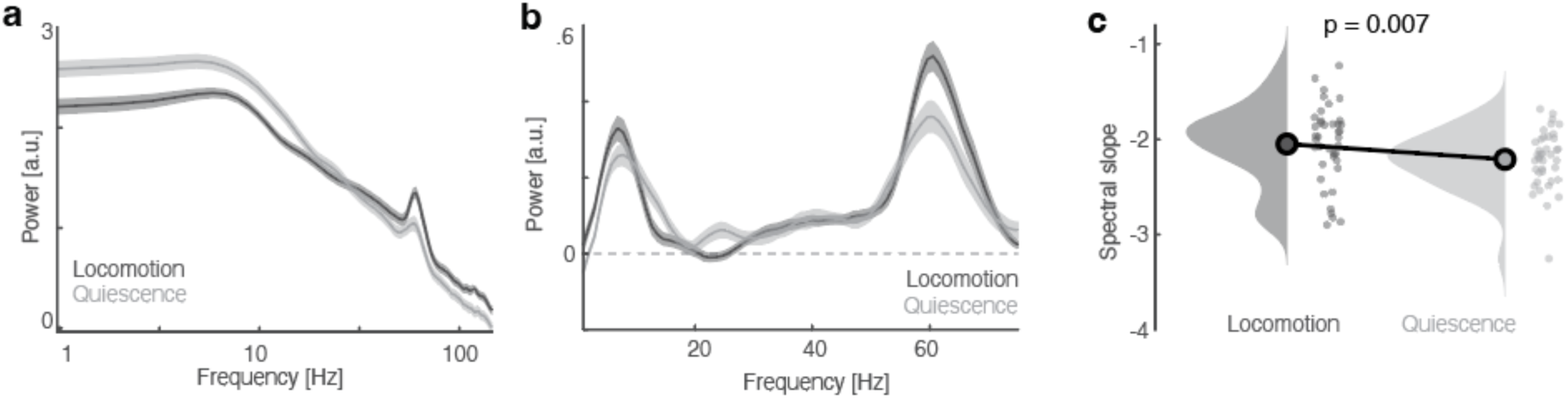
The spectral tilt during locomotion is independent of evoked gamma activity. (**a**) Grand-average power spectra during locomotion and quiescence without visual stimulation. (**b**) Oscillatory residuals. Note the prominent absence of the visual gamma peak around ∼20-40 Hz. (**c**) The spectral slope increased during locomotion (-2.05 ± 0.07) as compared to quiescence (-2.21 ± 0.05), which was statistically significant (p = 0.0070, t39 = 2.85, paired t-test). Related to Figure 2.

**Supplementary Fig. S3.**
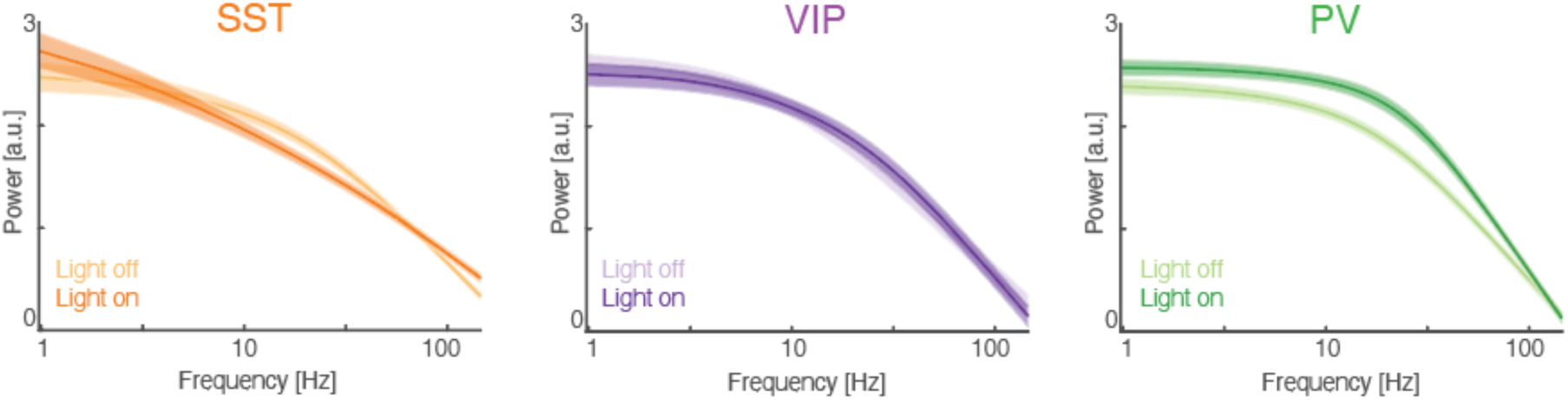
Aperiodic component of the power spectrum. Aperiodic background fits for SST (left), VIP (center) and PV (right) groups, highlighting the differences in background activity across the different groups. Related to Figure 5.

**Supplementary Fig. S4.**
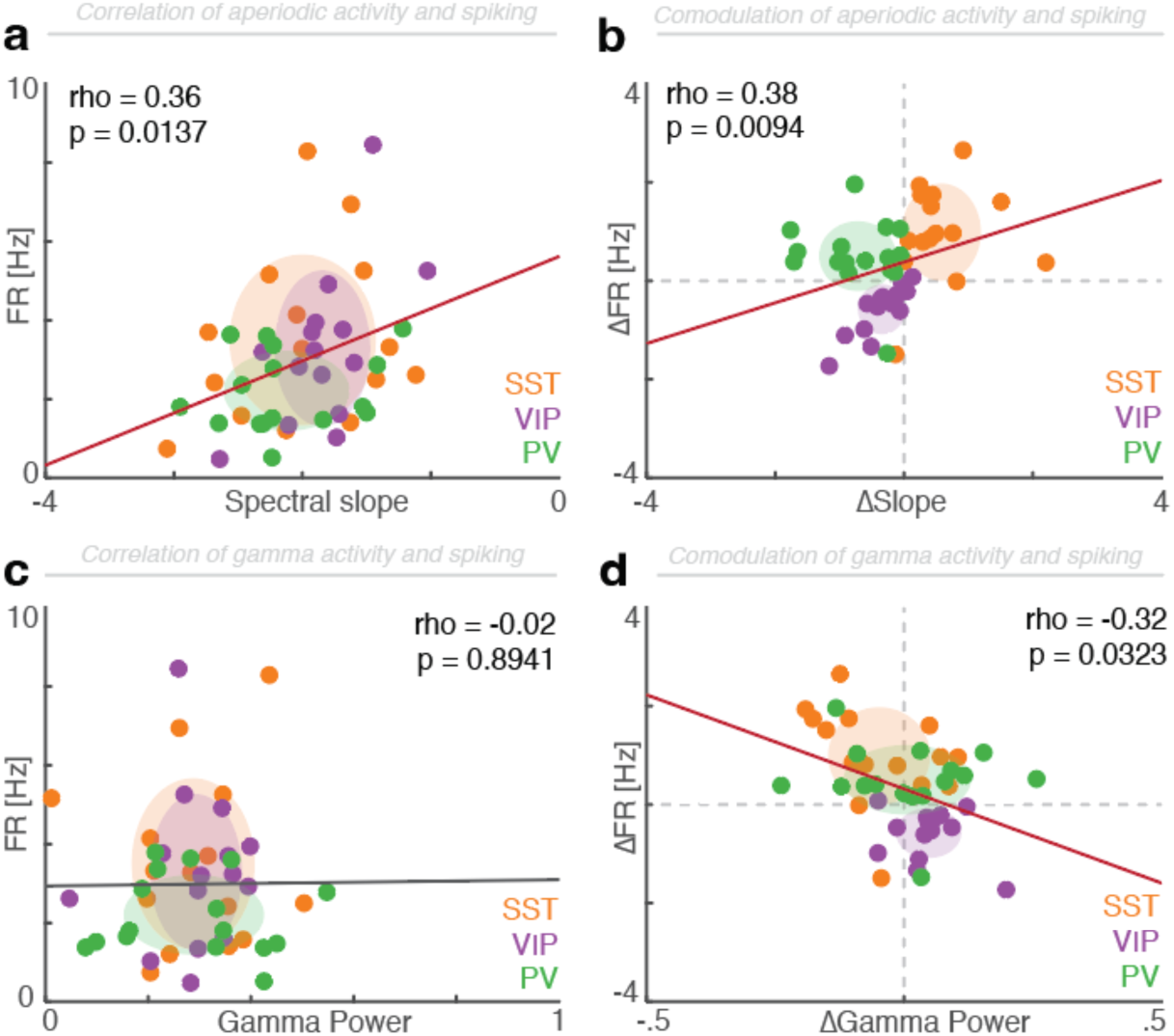
The relationship of neural spiking, oscillatory and aperiodic activity when averaged across all visual stimulation conditions. (**a**) Statistically significant correlation between the spectral slope and pyramidal cell spiking (p = 0.0137, rho = 0.36, Spearman correlation). Flatter spectral slopes signal increased neural spiking. (**b**) Optogenetic modulation (light on *minus* light off) demonstrates a significant relationship with a clear differentiation and compartmentalization of the different interneuron cell types; replicating the main effect (p = 0.0094, rho = 0.38, Spearman correlation). (**c**) Visual gamma power was not correlated with the overall spiking rate when all visual stimulation conditions were averaged (p = 0.8941, rho = -0.02, Spearman correlation). (**d**) However, the optogenetically-induced gamma power modulation (light on *minus* light off) demonstrated a significant negative correlation highlighting that elevated visual gamma synchrony predicted lower spiking rates (p = 0.0323, rho = -0.32, Spearman correlation). Related to Figure 6.

**Supplementary Fig. S5.**
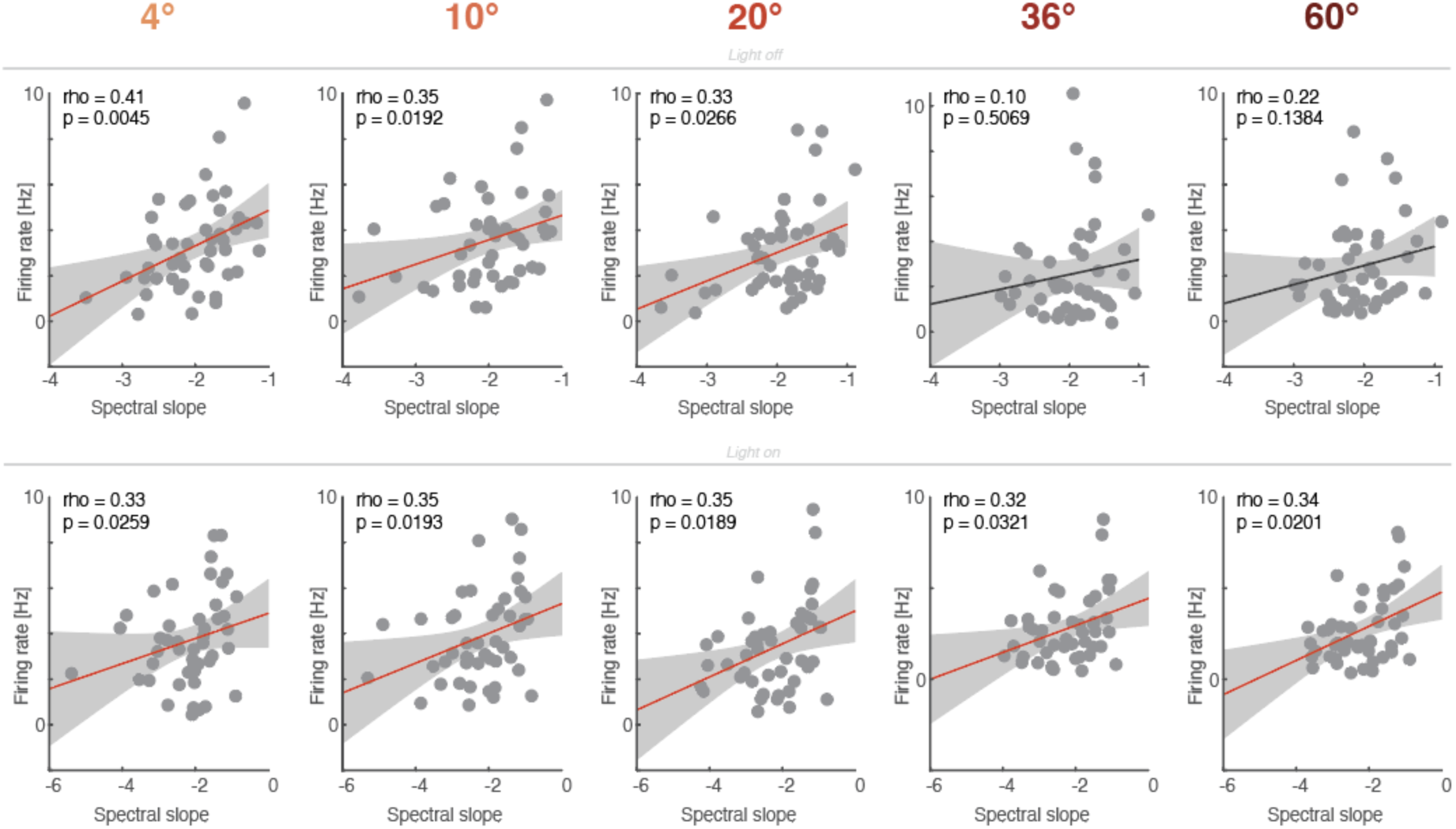
Correlation between the aperiodic slope and spiking activity as a function of different visual stimulus sizes. Left to right: Correlation between the spectral slope and neural firing rate (pyramidal cells) as a function of different stimulus sizes, which give rise to varying levels of gamma activity (cf. Figure 4). *Top row*: During the light off condition. Uncorrected spearman rank correlation coefficients are depicted (red regression lines indicate uncorrected p < 0.05). Same data as in Figure 6f, but not squared. *Bottom row*: During the light on condition. Same conventions as in the top row.

**Supplementary Fig. S6.**
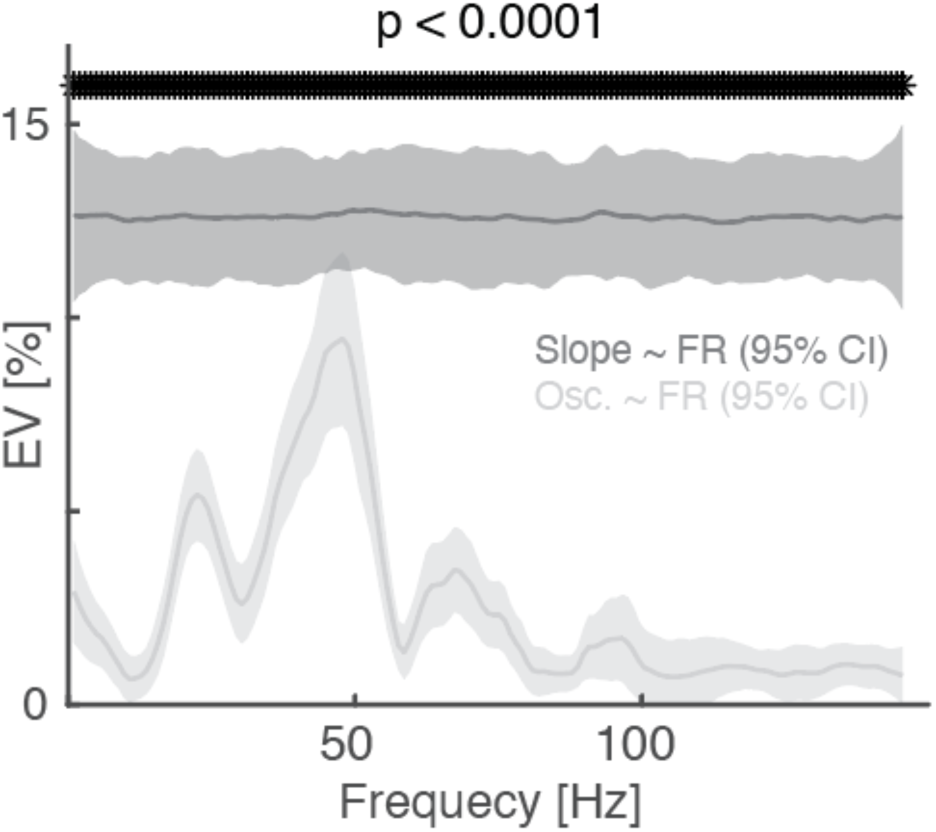
Explained variance of the spiking rate as a function of aperiodic (dark grey) and oscillatory activity. Explained variance (squared correlation coefficient) of the firing rate (FR) as a function of the aperiodic slope (dark grey) and oscillatory activity (light grey; bootstrapped 95% confidence intervals across 100 iterations that subsampled 90% of the subjects; smoothed with a 10 Hz window for visualization). A cluster-based permutation test revealed a large cluster that spanned all frequencies (p < 0.0001), thus, indicating that aperiodic activity explained more variance than oscillatory activity, irrespective of the peak frequency. Note that the visual gamma activity (∼20-40 Hz) also explained some variance (but note the negative association between the spiking and gamma modulation; cf. Figure 6d and **Figure S4d**), underscoring that a high-degree of synchrony may govern the relationship between neural spiking and aperiodic activity. Light-off condition, averaged across all visual stimulation conditions during locomotion. Aperiodic activity explained more variance about neural spiking than oscillatory activity. Related to Figure 6.

